# Hot spots drive uptake and short-term processing of organic and inorganic carbon and nitrogen in intertidal sediments

**DOI:** 10.1101/2023.11.29.569196

**Authors:** Philip M. Riekenberg, Bradley D. Eyre, Marcel T.J. van der Meer, Joanne M. Oakes

**Author notes:** Author contribution: PMR: Conceptualization (lead); writing-original draft (lead); formal analysis (lead); BDE: writing-review and editing (equal); funding acquisition (equal); formal analysis (supporting) MTJvdM: supervision (equal); writing-review and editing (equal); JMO: funding acquisition (equal); supervision (equal); writing-original draft (supporting); writing review and editing (equal).

## Abstract

This study uses dual-labelled (^13^C and ^15^N) stable isotope applications to examine uptake and short-term processing of carbon (C) and nitrogen (N) by microbial communities in intertidal sediment from three subtropical estuarine sites. We examine differences in microbial uptake and retention that arise due to domination of microbial processing by either microphytobenthos or heterotrophic bacteria. We compare amino acids and algal dissolved organic matter (Algal DOM) and glucose and NH_4_^+^ versus newly fixed microphytobenthos C (MPB-C) and NH_4_^+^ using *in situ* applications across 24 h to identify uptake into the microbial community and sediment OM. Algal DOM had preferential C uptake and more retention across 24 h indicating precursors incorporated into biosynthetic pathways for biomass. Conversely, amino acid C was not incorporated or rapidly respired to DIC but displayed clear preferential uptake and retention of ^15^N. Short-term (24 h) retention of glucose was higher than MPB-C, while uptake of ^15^N from NH_4_^+^ was similar between treatments, potentially indicating glucose-stimulated export of ^15^N via coupled nitrification-dentrification. Despite careful selection of similar sites and sediment types, we found substantial variability between replicates and sites in the uptake and processing of labeled substrate that challenged traditional statistical analysis due to non-homogenous variance. Uptake variability across orders of magnitude is likely due to disproportionate processing of substrates occurring in hotspots of microbial processing within sediment. Development of analytical techniques to provide robust strategies to handle variability caused by abiotic and biotic factors will allow greater clarity surrounding *in situ* biogeochemical processing in intertidal environments.

## Introduction

Intertidal sediments are important sites for intercepting and processing organic and inorganic matter prior to its export to shallow coastal seas (Bauer et al., 2013). Within euphotic intertidal sediments microbial communities dominated by microphytobenthos (diatom and cyanobacteria) are important mediators for both carbon (C) and nitrogen (N) derived from inorganic and organic substrates (Middelburg et al., 2000). Fixation of inorganic C via primary production coincides with high uptake and competition for N by autotrophs that can limit N for heterotrophs, depending on nutrient availability (Cook et al., 2007; Evrard et al., 2012; Riekenberg et al., 2020). Sediment C and N uptake and processing pathways have been relatively well characterized using stable isotope labelling techniques, (see below), but there is less evidence for how substrate quality, and small-scale variability in uptake influences these processes. Previous work has often had minimal replication primarily due to cost and the labour-intensive processing for samples supporting this work despite finding considerable differences between uptake rates for ^13^C and ^15^N within a variety of single site intertidal studies (coefficient of variation (%CV); Table 1).

**Table 1:**
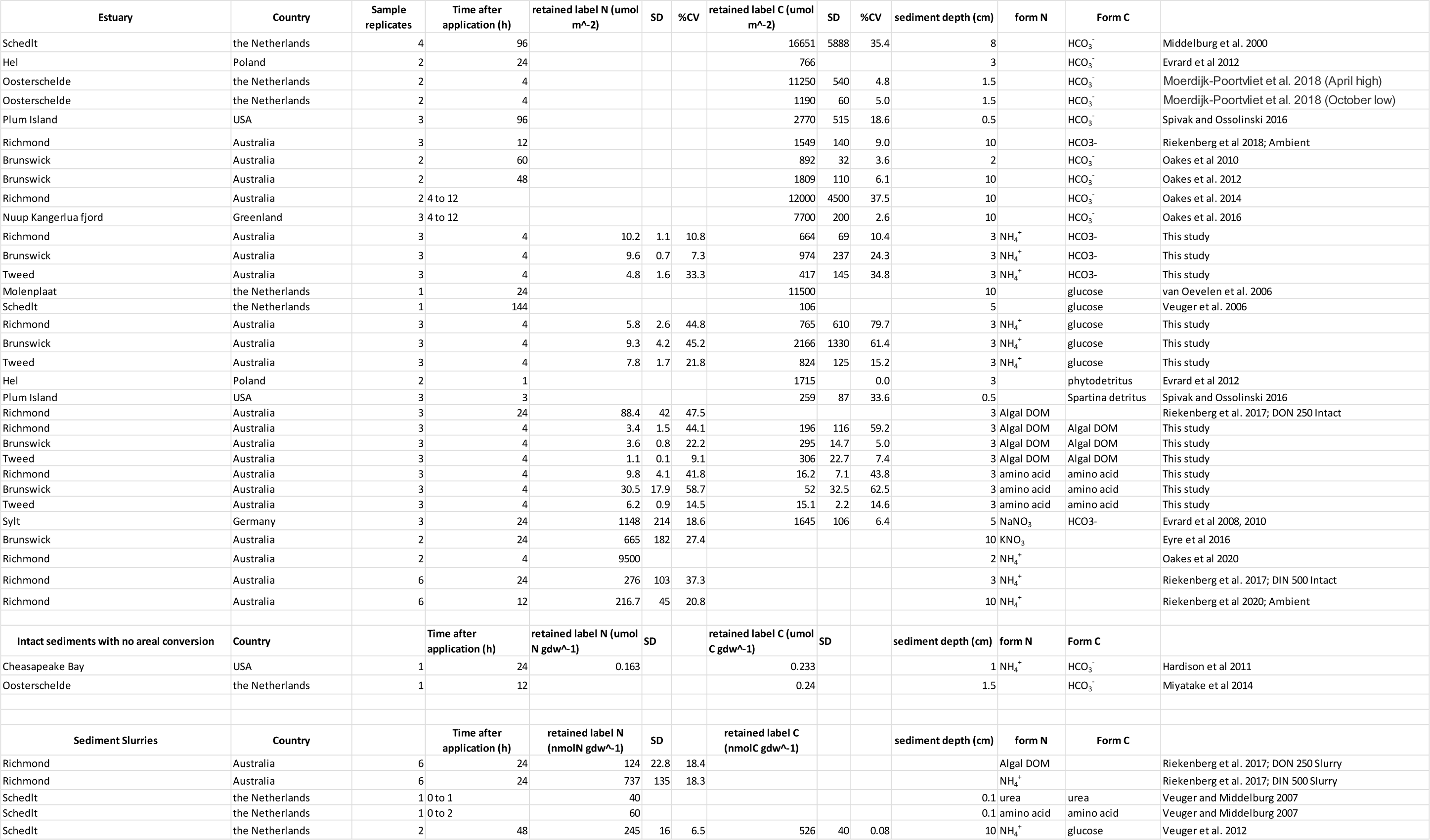
Compilation of studies examining uptake of ^13^C and/or ^15^N from organic or inorganic substrates into bulk measured intertidal sediments. SD indicates standard deviation where applicable.

Short term inorganic C uptake and processing is dominated by quick uptake and short-term retention of C in MPB biomass (Moerdijk-Poortvliet et al., 2018) followed by longer term retention of microphytobenthos derived carbon (MPB-C) in the sediments (30 d+) as extracellular polymeric substances (EPS) and reworked detrital MPB biomass (Oakes and Eyre, 2014; Oakes et al., 2013; Riekenberg et al., 2018). Newly fixed MPB-C is primarily exported via respiration and efflux of dissolved inorganic carbon (DIC) to the water column. MPB-C that is retained over longer periods is generally transferred to heterotrophic microbes and higher consumers via MPB exudates, which comprise a snot-like complex pool of substrates MPB exudates are sugar rich (primarily glucose) with labile components that are quickly (<24 h) used by heterotrophic bacteria (Oakes et al., 2012). More refractory components (e.g., carbohydrates in EPS) are processed more slowly (>24 h) as they require enzymatic break down into simpler components prior to use by heterotrophic bacteria (Miyatake et al., 2014). EPS can account for a considerable (29%; Cook et al. 2007) to marginal (10%; Oakes et al. 2010) portion of sediment organic carbon that can have implications for the function and support of intertidal food webs (Christianen et al., 2017; Riekenberg et al., 2022). Fixation of carbon, and production of EPS, is stimulated under high light environments (Goto et al., 1999; McMinn and Lee, 2018) as diatoms use EPS to make their surrounding environmental conditions more favorable. EPS helps to protect diatoms from excessive UV exposure, whilst also stabilizing sediments (minimizing diatom resuspension), and facilitating migration by motile diatoms (Hoagland et al., 1993).

Short term nitrogen uptake and processing via MPB-mediated pathways are less well characterized, but tend to be dominated by short term uptake of inorganic and organic materials from the overlying water column (Eyre et al., 2016a; Veuger and Middelburg, 2007) followed by limited transfer of N to heterotrophic bacteria (Evrard et al., 2008). In nutrient limiting settings MPB can outcompete heterotrophic bacteria for nitrogen substrates. N limitation of the microbial community results in strong coupling between MPB and heterotrophic bacteria that causes efficient recycling and strong retention of N at the sediment-water interface (Cook et al., 2007; Risgaard-Petersen et al., 2004). Increased nitrogen availability from the overlying water column makes the coupling between heterotrophic bacteria and MPB less efficient (Oakes et al., 2020), causing an increased accumulation of newly fixed N into the uncharacterized sediment pool (Riekenberg et al., 2020). Intertidal uptake of N is dominated by incorporation into the microbial community followed by loss of N as N_2_ via coupled nitrification-denitrification (Eyre et al., 2016b) or via physical loss (Oakes et al., 2020). Denitrification can be stimulated by access to a labile C source (Oakes et al., 2011) and is a vital pathway for nitrogen removal in estuarine sediments (Slater and Capone, 1987) that varies considerably within intertidal and subtidal ranges of subtropical estuaries (Eyre et al., 2016a; Oakes et al., 2020) and coastal shelf seas (Chua et al. 2021), ranging from a minimal to considerable pathway for N export (<1 to >100 µmolN_2_m^-2^h^-^1; Douglas et al., 2022).

An array of potential C and N substrates are available to microbial communities ranging in complexity from glucose and ammonium to lignin (Bronk et al., 2007; See et al., 2006). However, substrate bioavailability and oxygen availability largely determine whether these sources are used by microbes (Berman and Bronk, 2003). Substrate use by the microbial community varies as uptake can be limited by secondary limitations (e.g. C, N or phosphorous) or an inability to access the substrate through enzymatic action (Crawshaw et al., 2019; Riekenberg et al., 2017). Thus, substrate availability does not necessarily indicate microbial usage. The net role of oxygen, enzyme, and secondary nutrient availability will influence benthic N metabolism, especially when organic matter is the predominate source supporting the microbial community. The stoichiometry (C:N) and complexity of organic matter govern microbial uptake, with the relative lability of OM contributing to denitrification efficiency within systems (Albert et al., 2021; Eyre et al., 2013; Mayali et al., 2013; Oakes et al., 2011). The complexity of carbon substrates can determine which heterotrophs thrive within benthic microbial communities (Abell et al., 2013; Carlson et al., 2020) and marine biomass is an important source of dissolved proteins and peptides supporting heterotrophic bacteria in the intertidal zone (Schmidt et al., 2017; Seidel et al., 2015). Further examination of uptake and short-term processing using labeled organic and inorganic substrates will be useful in directly tracking substrate use and identifying rate differences in the processing pathways that potentially limit or drive benthic N metabolism (Eyre et al., 2016a; Oakes et al., 2020). The variability and relative efficiencies of uptake for carbon and nitrogen from organic and inorganic substrates within *in situ* intertidal sediments largely remain a knowledge gap. Application of combined isotope-labelled substrates may provide further insight into the short-term processing of organic matter and indicate which scenarios are likely to stimulate short-term denitrification (e.g. lability of OM, and competition between heterotrophs and microphytobenthos).

Here, we applied inorganic and organic substrates labelled with the rare stable isotopes ^13^C and ^15^N to in situ intertidal sediments to examine uptake and short-term retention (24 h) by the sediment microbial community within three subtropical estuaries. This work compares uptake and short-term processing across a range of bioavailable compounds (e.g. glucose to dissolved organic matter) with emphasis on 1) comparing the microbial processing of carbon and nitrogen from organic versus inorganic substrates 2) comparing the microbial uptake and processing of carbon and nitrogen from organic compounds across a range of lability and bioavailability and 3) quantifying the variability in substrate use both within and between subtropical estuaries from sites with similar characteristics. We expected 1) relatively labile substrates such as amino acids would have greater uptake and processing than more complex pools such as algal derived DON, 2) uptake of ^13^C and ^15^N to be dominated by MPB, and 3) the subsequent processing pathways for newly fixed OM to be dominated by support of primary production pathways.

## Methods

### Site description

The study was undertaken in the austral spring (August and September) of 2017 in three river-dominated subtropical estuaries: the Richmond, Tweed and Brunswick River estuaries in New South Wales, Australia (Fig. 1A). Across the study period these sites had a semidiurnal tidal range of ∼2 m with site water salinity ranging from 25.2 to 28.6. This range is typical of Australian subtropical rivers sampled close to the river mouth during low flow conditions which occur in the winter/spring transition (Ferguson et al., 2004). Water column dissolved inorganic nitrogen (DIN) values typical of this period are 1.4, 1.9, and 2.4 µmol L^-1^ with DON values of 8.3, 9.6 and 11.8 µmol L^-1^ for the Brunswick, Tweed and Richmond, respectively (Eyre, 2000), with DON comprising the largest portion of total nitrogen, which is typical in subtropical estuaries during low flow conditions (McKee et al., 2000). Flow rates across the year can be considerably larger and quite variable due to episodic flushing usually associated with low pressure systems or tropical cyclones during the wet season summer/autumn (Ferguson et al., 2004) and this variability in flow can impact sediment processing (Oakes et al. 2020; Oakes and Eyre 2014).

**Figure 1:**
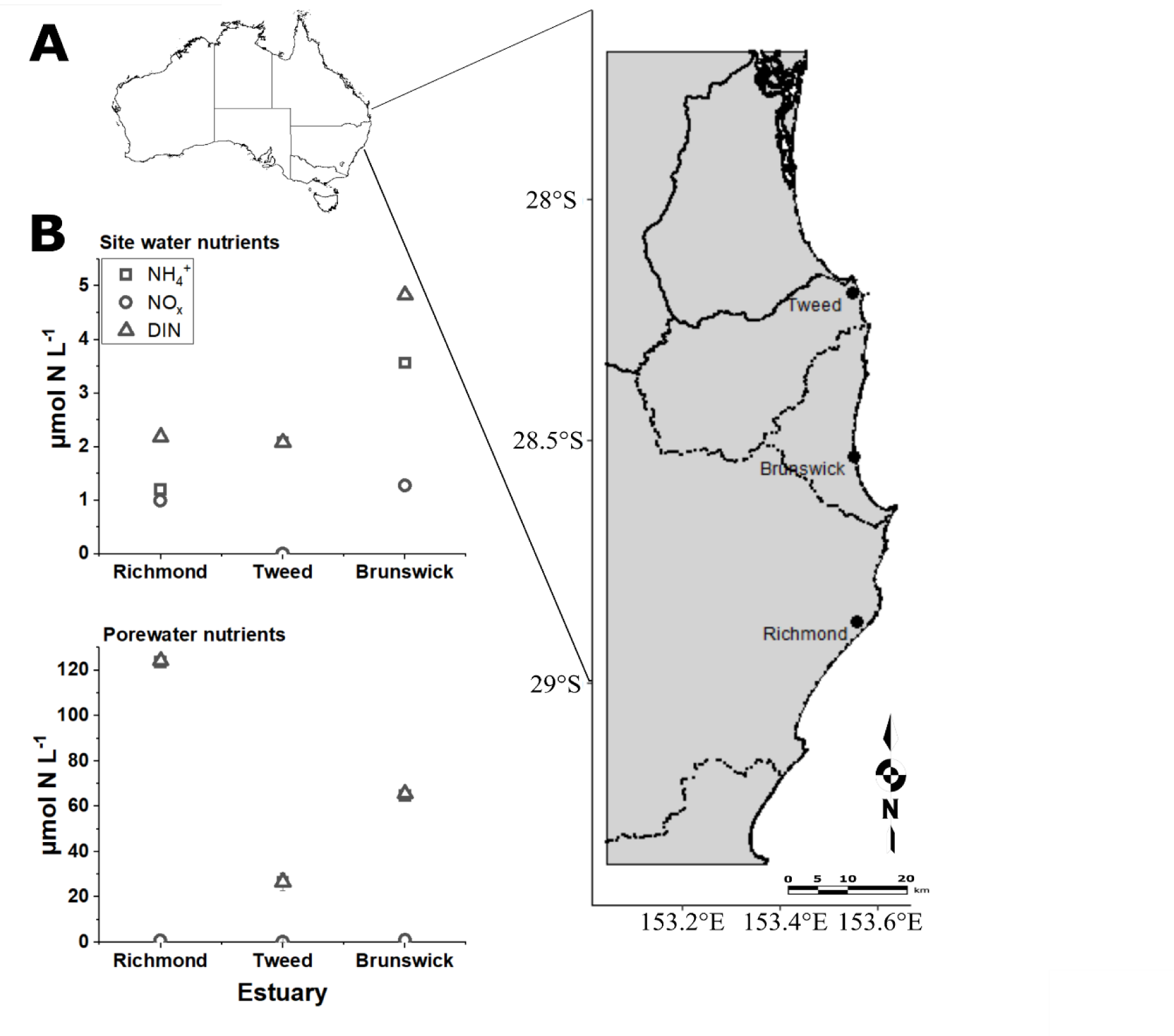
A) Study site locations in New South Wales, Australia in the Tweed, Brunswick and Richmond River catchments (black outlines) and B) site water and porewater NH_4_^+^, NO_x_, and DIN (NH_4_^+^ + NO_x_) values. Sampled mudflats were within 2 km straight line of the river mouths to the Pacific Ocean within each estuary.

### 13C and ^15^N labeling additions

There are four treatment applications in this study: 1) algal DOM extracted from diatoms raised on labelled NaHCO_3_ and NH_4_ ^+^ resulting in atom% values of 37.1 for ^15^N and 5 for ^13^C, 2) an amino acid mixture labelled 99 atom% for both ^13^C and ^15^N, 3) glucose and NH ^+^, both labelled 99 atom% for ^13^C and ^15^N, respectively, and 4) NaHCO_3_ and NH_4_ ^+^ labelled 98 and 99 atom% for ^13^C and ^15^N, respectively. Further detail about preparation and concentration of each treatment is described below.

The diatom-derived dissolved organic matter treatment (algal DOM; treatment 1) was prepared following the method in Riekenberg et al. (2017) using an axenic culture of the diatom *Thalassiosira pseudonana* grown in media containing 99% ^15^NH_4_^+^ and 98% NaH^13^CO_3_ (Cambridge Isotope Laboratories). The extracted DOM had a ^15^N atom% of 37.1 and a ^13^C atom% of 5 (corresponding to ∼160,000‰ and 3,800‰, respectively). A small concentration of remaining DIN represented 2% of the total N in the treatment as measured via flow injection analysis after dilution and persulfate digestion (Lachat, 1994). Any remaining dissolved inorganic carbon was removed through acidification with 2N HCl prior to filtration of algal extract through a 0.45 µm syringe filter (Minisart, Sartorius). The diatom-derived DOM was divided into aliquots prior to freeze drying that were determined to be sufficient to produce a target concentration of 60 µmol L^-1^ for DON and 162 µmol L^-1^ for DOC when resuspended with 20 ml of filtered site water (0.2 µm, Sartorius).

The amino acid treatment (amino acid mixture; treatment 2) was prepared from a commercially available 99 atom% dual-labelled (^13^C and ^15^N) mixture of 20 amino acids (aspartic acid, glutamic acid, asparagine, serine, glutamine, histidine, glycine, threonine, alanine, arginine, tyrosine, cystine, valine, methionine, tryptophan, phenylalanine, isoleucine, lysine, and proline; Cambridge Isotope Laboratories). The amino acid mixture was suspended in deionized water and divided into aliquots prior to freeze drying that were determined to to be sufficient to produce a target concentration of 60 µmol L^-1^ for DON and 211 µmol L^-1^ for DOC when resuspended with 20 ml of filtered site water (0.2 µm, Sartorius).

Mixtures of glucose and NH_4_^+^ (treatment 3) and NaHCO_3_ and NH_4_^+^ (treatment 4) mixtures were prepared from 99 atom% ^13^C-labelled glucose, 99 atom% ^15^N-labelled NH_4_^+^, and 98 atom% ^13^C-labeled NaHCO_3_. These two treatments were suspended in deionized water and divided into aliquots prior to freeze drying that were determined to be sufficient to produce a target concentration of 60 and 20 µmol N L^-1^ and 1100 and 1800 µmol C L^-1^, respectively, when resuspended with 20 ml of filtered site water (0.2 µm, Sartorius).

### Labeled substrate application

Treatments were applied in August of 2017 (austral spring) to intertidal sediment at sites within 2 km straight line distance upstream from the river mouths of the Richmond (28° 52’ 30” S, 153° 33’ 26” E), Tweed (28° 11’ 43.1″ S 153° 32’ 49.2″ E), and Brunswick (28° 32’ 01.8″ S 153° 33’ 07.1″ E) Rivers in New South Wales, Australia (Fig. 1A). Particular care was taken to select sites that were similar within each estuary (i.e., similar location within estuary, sandy mud, similar inundation times) to minimize environmental variability between sites. Areas within sites that were free of animal burrows or wallows were selected to minimize 1) transfer of label between plots due to foraging, 2) channeling of initial treatment application to deeper layers, or 3) uneven resuspension and flushing of sediments during inundation due to wallows. At each site, four quadrats (one per treatment; 60 × 60 cm) were placed 2 m apart from one another at the same height within the intertidal zone. Quadrats were divided into 9 plots (20 × 20 cm) to allow for 4 replicate samplings for 4 h and 24 h periods for each treatment and leaving one blank per quadrat. Quadrats were anchored in two corners by 20 cm lengths of PVC pipe pushed into the mud flat that remained in place between samplings to mark the quadrat positions. Two loggers for temperature and light (HOBO) were attached to anchoring posts on opposite corners of the outside quadrats across each site.

Treatments were applied to bare sediment on a falling tide during mid-day sunlight conditions to ensure maximum time (1.5-4 h) for uptake of treatment applications prior to tidal inundation flushing the majority of the unincorporated application from the sediment. Each application was sprayed onto the sediment using mechanical sprayers to evenly disperse 20 ml of treatment onto each of 8 plots within the treatment quadrat. Labeling dosages ranged from 10 to 113 µmol C m^-2^ and 0.1 to 1.9 µmol N m^-2^ (supplemental Fig. 1) amongst the treatments. Concentrations were higher than those naturally occurring to provide a pulse of labeled material that would remain detectable after tidal flushing. Treatment applications were applied individually to avoid any cross contamination between treatments. Within treatment quadrats, individual plots were randomly allocated to 4 h and 24 h samplings with 4 replicates taken for each sampling period. This sampling design has been previously found to be statistically independent (Riekenberg et al. 2017) and does not represent a reduction of variance between samplings (pseudoreplication) due to the high variability found in sediment at the millimeter to centimeter scale. The majority of unincorporated labelled material was removed from the sediment due to replacement and turnover of porewater during tidal inundation prior to sampling. Removal of both ^13^C and ^15^N at 4 h was confirmed by the highest remaining δ^13^C and δ^15^N values left in sediment organic matter being 194‰ and 591‰, respectively, which reflect a small fraction (1.3% and 0.6%) of the initial treatment remaining. Any remaining, but unincorporated material from treatment applications was found to be minimal (26±50‰ and 7±25‰ for δ^15^N and δ^13^C, respectively) through comparison between raw and KCl-extracted (for ^15^N) or acidified (for ^13^C) sediments from the 0-1 cm layer at 24 h.

### Sample collection

Prior to treatment application four cores (9 cm diameter × 3cm depth) were sampled from between the quadrats and sectioned into 1 cm depths to 3 cm, including a 1 mm top scrape and a 1cm^3^ sample for chlorophyll-*a* analysis, serving as controls for sediment characterization and δ^13^C and δ^15^N values. All sections were placed in individual bags and stored frozen (-20°C), while the 0-1 cm depth was immersed in liquid nitrogen and flash frozen directly to ensure preservation of the microbial community for biomarker analysis prior to transport back to the laboratory. Within the quadrats, at 4 h and 24 h after the treatment applications, four cores were sampled, similarly sectioned, and stored frozen until freeze drying occurred within the laboratory. Prior to label applications, water from site porewaters and the overlying water column was collected and passed through a 0.7µm syringe filter and frozen prior to analysis for baseline nutrient concentrations (Lachat, 1994) and δ^15^N values for NO_3_^-^ via the bacterial denitrifier method (Sigman et al., 2001).

### Sample analysis

Chlorophyll *a* was measured by colorimetry (Lorenzen, 1967) within the 0-1 cm depth for both control and treatment application cores at each site. MPB-C biomass was calculated using a conversion factor of 40 for C:chlorophyll-*a* which is in the range reported for microalgae in subtropical Australian estuaries (30-60; (Ferguson et al., 2004; Riekenberg et al., 2018). MPB-C biomass estimated in this way was only used as a measure to compare biomass between estuaries and treatment applications and was not used for subsequent determination of ^13^C or ^15^N uptake or retention.

Frozen sediment was freeze dried, homogenized, and a 3 cm^3^ subsample was extracted with (2 M KCl indundation and centrifugation to remove 3× followed with 3× inundation and centrifugation with double distilled water), or a 20 to 40 mg subsample acidified (two drop 2N HCl into silver cups) to remove any adsorbed inorganic and organic N and inorganic C, respectively. Washed or acidified sediment was dried (60°C) and weighed into tin or silver cups for analysis of %N and δ^15^N values or %C and δ^13^C values, respectively, using a Flash 200 elemental analyzer coupled to a Delta V Advantage isotope ratio mass spectrometer (Thermo Scientific, Bremen). Reproducibility of δ^13^C and δ^15^N values was ±0.1‰ and ±0.2‰ for samples with enrichment less than 100‰ and decreased for enrichment beyond that range.

To partition the uptake of ^15^N into microphytobenthos and heterotrophic bacteria, we used amino acid analysis of alanine to determine the relative uptake of ^15^N between the D- and L-forms. A subsample of sediment from the 0-1 cm depth (3 g) was acidified (2N HCl, 15 minutes or until bubbling stopped) to remove carbonates, hydrolyzed overnight in 6N HCl at 110°C, purified using cation exchange chromatography (Dowex 50WX8-100), and derivatized into N-pivaloyl-amino acid-i-propyl esters. The concentrations and δ^15^N values of the N contained in the chiral forms of alanine (D/L-Ala) were examined via gas chromatography-combustion isotope ratio mass spectrometer using 50 m of CP-Chirasil Val columns (2 25 m × 0.25 mm ×0.12 µm, Agilent) mounted in an Agilent 6890 gas chromatograph connected to a Delta V advantage isotope ratio mass spectrometer via a combustion III interface. Daily check standards were performed using an in-house standard mix of D- and L-alanine and norleucine to confirm adequate separation between the chiral forms and to monitor the reference standard (norleucine) δ^15^N values to ensure adequate oxidation of the reactor across runs. Samples for D/L-Ala were only examined from the 0-1 cm depth due to previous work having confirmed that most ^15^N uptake is confined to the 0-1 cm depth (>85%; Riekenberg et al. 2017) and limitations of time and cost associated with the laborious nature of this analysis.

### Calculations

^13^C and ^15^N data are presented as excess µmol ^13^C or ^15^N per square meter of sediment to allow for direct comparison between estuarine sediments with different porosities, calculated as:

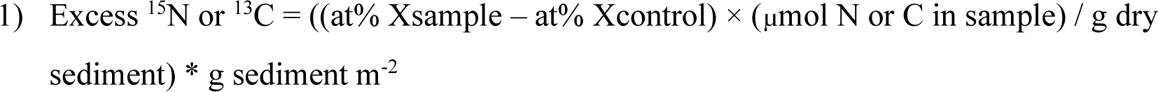

where at% X is the percentage of C or N atoms present as ^13^C or ^15^N in the sample or control sediment, and g sediment m^-2^ is the dry weight of sediment remaining when porosity is accounted for in each estuarine sediment. To account for the difference in atom % for both ^13^C and ^15^N in the algal DOM treatment and the other treatments, we adjusted the values by 20× and 2.7×, respectively. To provide ^13^C/^15^N ratios, the sum of excess ^13^C and ^15^N was calculated across the 0-3 cm sediment depths examined.

#### Statistical analysis

Variance for ^13^C and ^15^N areal uptake and ^13^C/^15^N ratios was significantly different (Levene’s HOV, *p*<0.05) for each of the variables (treatment, estuary, sampling) time regardless of the transformations indicated by the bestNormalize package in R (Peterson and Cavanaugh, 2020). Therefore, comparisons between organic and inorganic treatments among estuaries were made using non-parametric pairwise Wilcoxon sign-rank tests. ANOVAs were used to compare biomass contributions for heterotrophic bacteria, where variance was homogenous (Levene’s F_2,69_=1.8, *p*=0.2). Biomass contributions were similar between the 4 and 24 hour periods (3-way ANOVA: F_1,71_=2.6, *p*=0.1, Hour) with no significant interaction between Hour and Estuary or Treatment. We therefore proceeded with 2-Way ANOVAs for Estuary*Treatment. For the contribution of heterotrophic bacteria (HB%) from D/L-Ala measurements we found a significant 3-way interaction (3-way ANOVA: F_6,70_=2.3, *p*=0.05, Estuary*Treatment*Hour). Variance was homogenous within the data set (Levene’s F_3,67_=1.52, *p*=0.2) and Hour was not significantly different (*p=*0.06) as a variable or in interactions with Estuary and Treatment so we performed separate 2-Way ANOVAs for Estuary*Treatment for the 4 and 24 h periods. To further examine the between site variability from this study and the variability observed across the wider literature, we used the R package cvequality (Version 0.1.3; Marwick and Krishnamoorthy 2019) to test for significant differences between the coefficient of variation for uptake rates for C and N substrates.

## Results

### Site characteristics

Surface water and porewater nutrients were dominated by NH_4_^+^ concentrations ranging from 2 to 4 µmol N L^-1^ and 27 to 124 µmol N L^-1^, respectively (Fig.1B), with NH_4_^+^ comprising >98% of porewater DIN and 54 -100% of surface water DIN. Water column and porewater concentrations for NO_3_^-^ were 0 to 1.2 µmol N L^-1^ with δ^15^N values of 6.6±10.5‰ and 29.8±2‰ across all three sites. Across all sites and depths, sediment was dominated by coarse and medium sand (75-90%), with mud fractions ranging from <0.5% in the Brunswick to 13% in the Tweed. Sediment density was higher and porosity was lower from the Tweed to the Brunswick, with the Richmond intermediate in both comparisons, ranging from 0.3 to 1.2 g m^-3^ and 0.9 to 0.3 g ml^-1^. Both sediment density and porosity varied significantly among estuaries (One-way ANOVA; density F_2,8_=45.7, *p*<0.001; porosity F_2,8_=125, *p*<0.001).

MPB-C biomass estimated from chlorophyll-*a* in the control cores ranged from 271±71 to 50±23 mmol C m^-2^ and was highest in the Richmond and comparably lower in the Brunswick and the Tweed estuaries (One-way ANOVA; F_2,17_=27.5, *p*<0.001). There was no significant difference between MPB-C biomass in control and treatment applications in the Richmond (Tukey’s HSD all *p*>0.05; One-way ANOVA, F_4,34_=6, *p*=0.001), the Brunswick (One-Way ANOVA, F_4,37_=1.3, *p*=0.3), or the Tweed estuaries (One-Way ANOVA, F_4,37_=2.7, *p*>0.05).

### Sediment uptake and short-term processing

Between the organic treatments (treatments 1 & 2), uptake of ^13^C m^-2^ was higher for algal DOM (treatment 1) than for the amino acid mixture (treatment 2) within all three estuaries (all *p*=0.036, Fig. 2A). Within each estuary there was no change in ^13^C m^-2^ (all *p*>0.05) from 4 to 24 hour for either treatment. For data pooled across estuaries, we again observed higher uptake of ^13^C m^-2^ for algal DOM than amino acids (*p*= 0.009 for both) at both 4 and 24 h (Fig. 2B) and no change in ^13^C uptake between 4 and 24 h (*p*>0.1). In contrast to ^13^C uptake in the organic treatments (1 & 2), uptake of ^15^N m^-2^ was higher from the amino acids than for algal DOM within all three estuaries (all *p*=0.036, Fig. 3A). Uptake of ^15^N m^-2^ was similar (all *p*>0.1) within each individual estuary for each treatment at 4 and 24 h after application. By pooling across estuaries, we found that uptake of ^15^N m^-2^ was higher for the amino acids treatment than for algal DOM at 24 h than at 4 h (Fig. 3B) and that uptake of algal DOM was higher at 24 h than at 4 h (*p*<0.01) but all other treatments had similar uptake (*p*>0.1) between the two time periods.

**Fig. 2:**
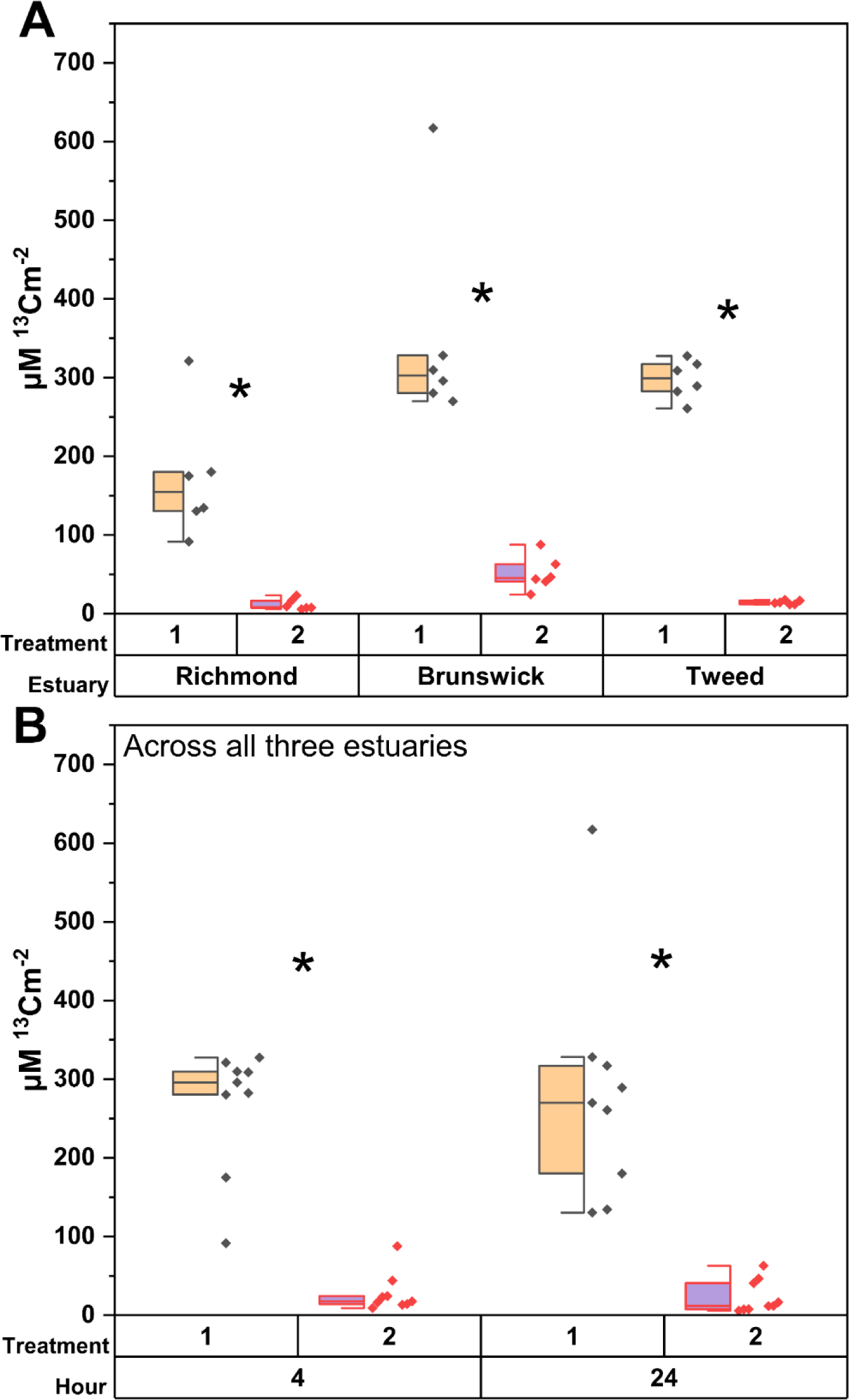
Uptake of ^13^C from organic treatment applications applied to intertidal sediment A) within each estuary and B) after 4 and 24 h across all three estuaries. Dual labelled (^15^N and ^13^C) treatment applications contain: 1) algal dissolved organic matter and 2) an amino acid mixture with uptake concentrations scaled to allow direct comparison between treatments. An asterisk indicates a significant difference (*p*<0.05) between pairings using non-parametric pairwise Wilcoxon sign rank tests.

**Figure 3:**
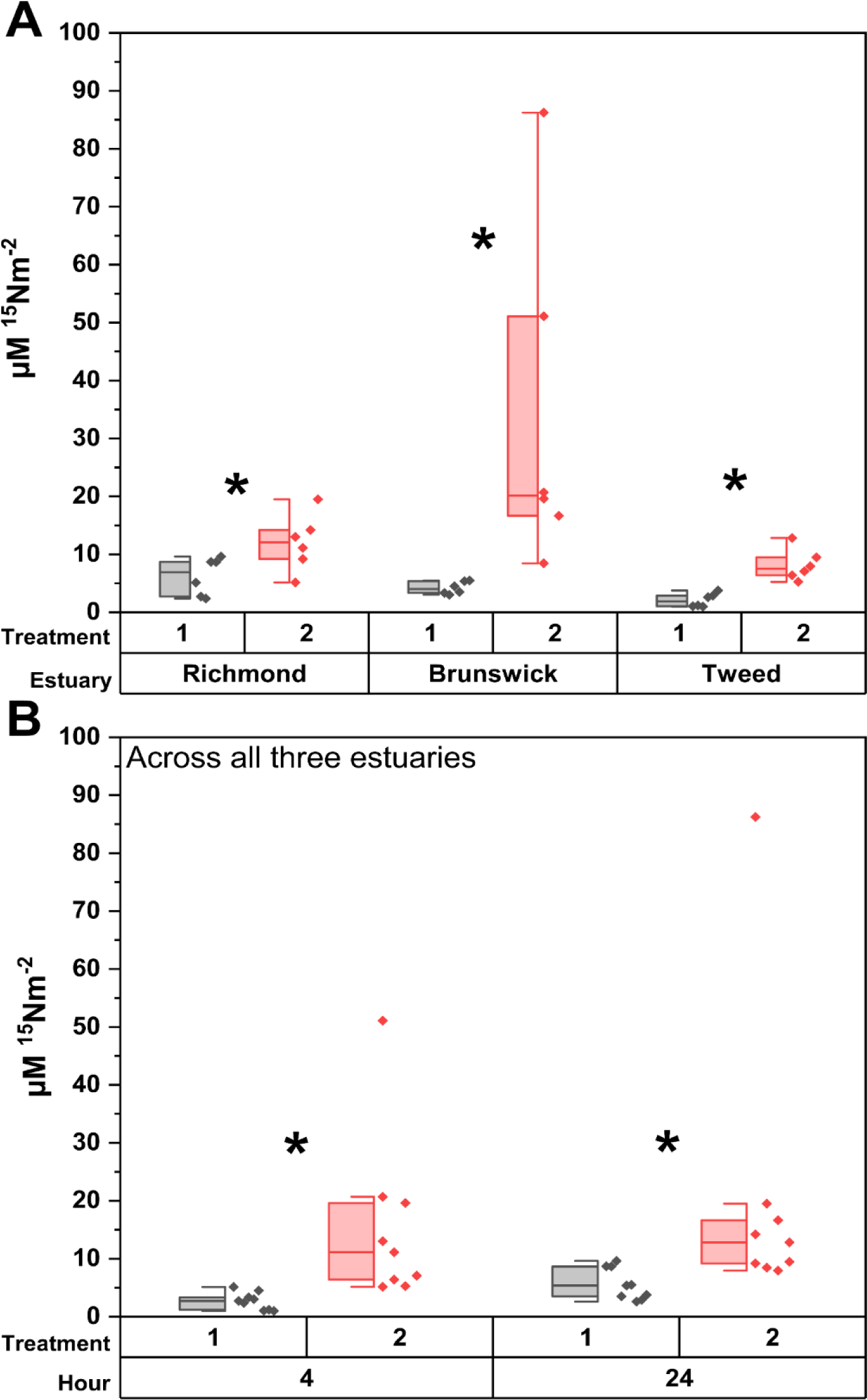
Uptake of ^15^N from treatment applications applied to intertidal sediment A) within each estuary and B) after 4 and 24 h across all three estuaries. Dual labelled (^15^N and ^13^C) treatment applications contain: 1) algal dissolved organic matter, and 2) an amino acid mixture. Uptake concentrations have been scaled to allow direct comparison between treatment applications. Asterisks indicate a significant difference (*p*<0.05) between pairings using non-parametric pairwise Wilcoxon sign rank tests.

Between the inorganic N treatments (treatment 3 & 4), uptake of ^13^C was higher for glucose than for NaHCO_3_^-^ in the Brunswick (*p*= 0.036), approached significantly higher in the Tweed (*p*=0.06), but did not differ in the Richmond River estuary (Fig. 4A). Within each estuary there was no change in ^13^C m^-2^ (all *p*>0.05) from 4 to 24 hour for either treatment. For data pooled across estuaries, we found that uptake of ^13^C m^-2^ was similar between treatments at 4 h but was higher for glucose than for NaHCO_3_^-^ at 24 h. Additionally, uptake of treatment 3 was nearly significantly higher at 4 h than at 24 h (Fig. 4B; *p=*0.06) but was similar across the two periods for treatment 3. Between treatments 3 and 4, uptake of ^15^N m^-2^ (Fig. 5A) was similar in the Brunswick, higher for treatment 4 in the Richmond (*p*= 0.036) and higher for treatment 3 in the Tweed (*p*=0.036). By pooling across estuaries, we found that uptake of ^15^N m^-2^ was similar between treatments 3 and 4 between estuaries (Fig. 5A; *p*>0.1) and time (Fig. 5B; *p*>0.1).

**Fig. 4:**
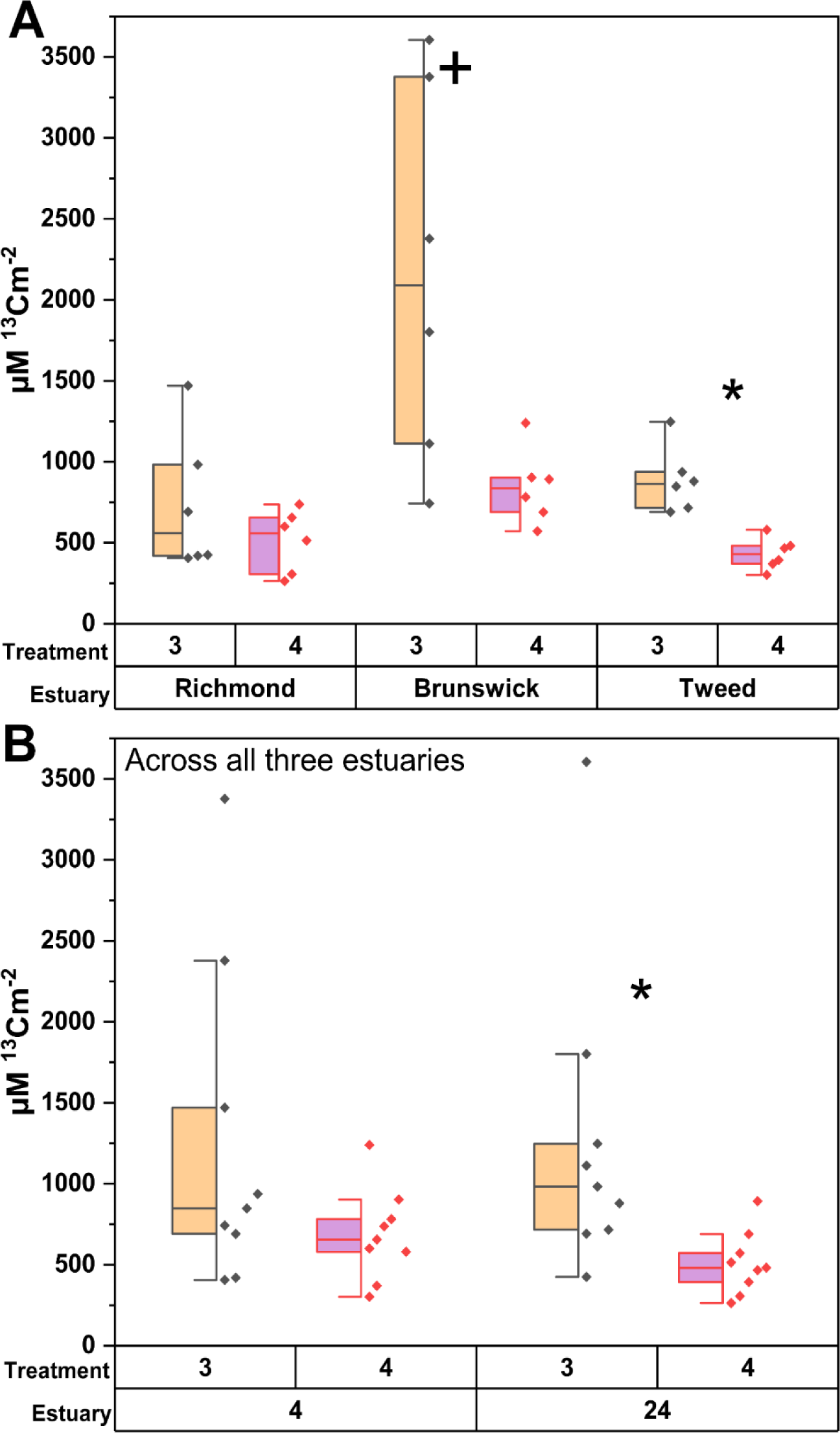
Uptake of ^13^C from inorganic treatment applications applied to intertidal sediment A) within each estuary and B) after 4 and 24 h across all three estuaries. Dual labelled (^15^N and ^13^C) treatment applications contain 3) glucose and NH_4_^+^, and 4) NaHCO_3_^-^ and NH_4_^+^ with uptake concentrations scaled to allow direct comparison between treatments. An asterisk indicates a significant difference (*p*<0.05) between pairings and the cross mark indicates a close to significant difference (p=0.06) using non-parametric pairwise Wilcoxon sign rank tests.

**Fig. 5:**
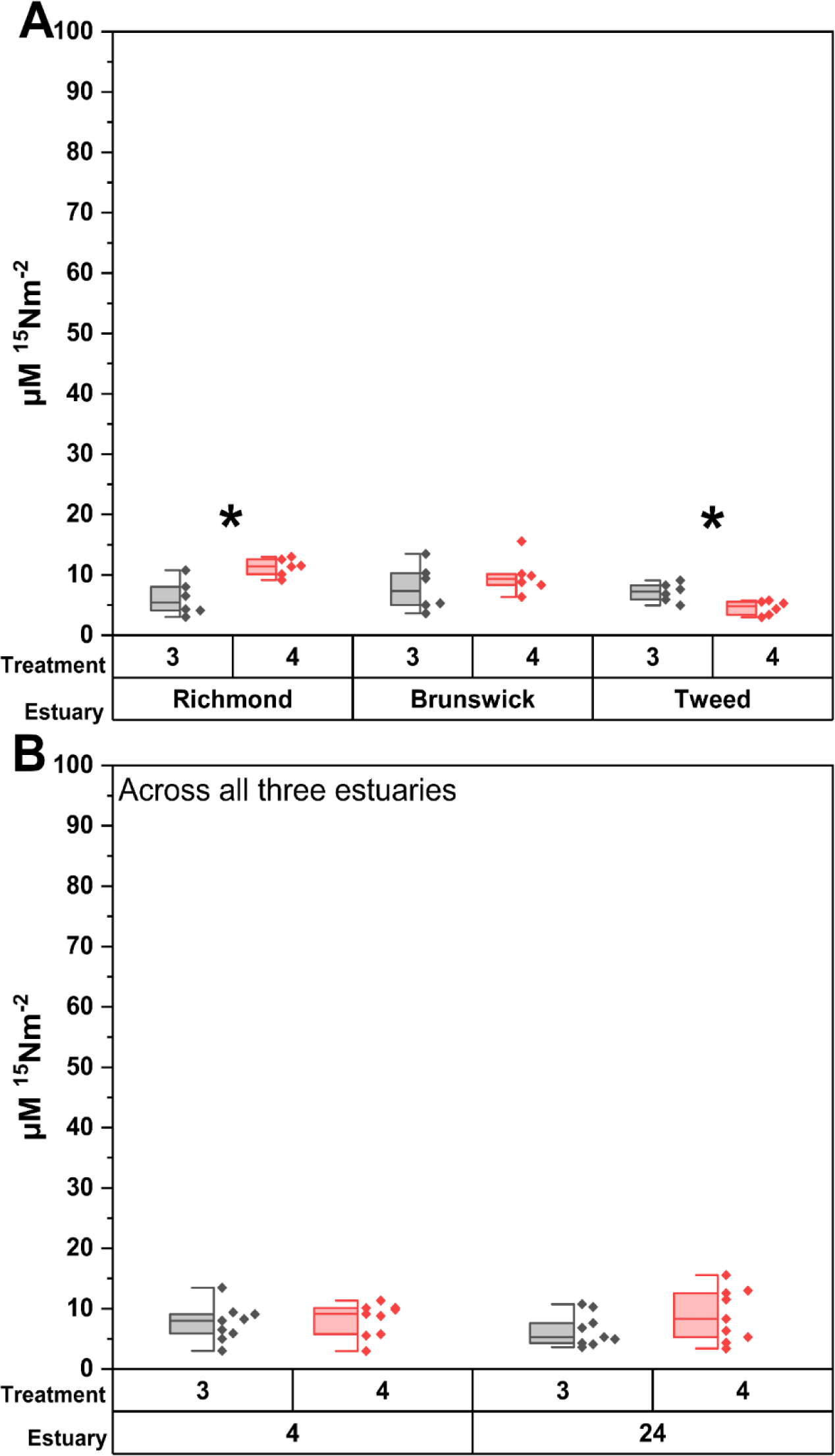
Uptake of ^15^N from inorganic treatment applications applied to intertidal sediment A) within each estuary and B) after 4 and 24 h across all three estuaries. Dual labelled (^15^N and ^13^C) treatment applications contain: 3) glucose and NH_4_^+^, and 4) NaHCO_3_ and NH_4_^+^ with uptake concentrations scaled to allow direct comparison between treatments. An asterisk indicates a significant difference (*p*<0.05) between pairings using non-parametric pairwise Wilcoxon sign rank tests.

^13^C/^15^N uptake ratios did not differ (all *p*>0.1) between 4 and 24 h samplings within each individual estuary. At all sites ^13^C/^15^N ratios were higher for algal DOM than for the amino acid treatment (treatment 1 vs 2; Fig. 6A; all *p*=0.036) but were only higher for the glucose/NH_4_^+^ treatment in the Tweed (Treatment 3 vs 4; Fig 6C; *p*=0.036). ^13^C/^15^N ratios were higher for the algal DOM treatment than for the amino acids at 4 and 24 h (treatment1 vs 2; Fig 6B; *p*= 0.009 for both) but were only higher for the glucose and NH_4_^+^ treatment at 24 h (treatment 3 vs 4; Fig. 6D; *p*=0.009). Comparing the coefficient of variation between the ^13^C (F_1,12_=19.4, *p*=0.047) and ^15^N (F_1,12_=13, *p*=0.29) applications indicated significantly different amounts of variance among the C substrate applications.

**Figure 6:**
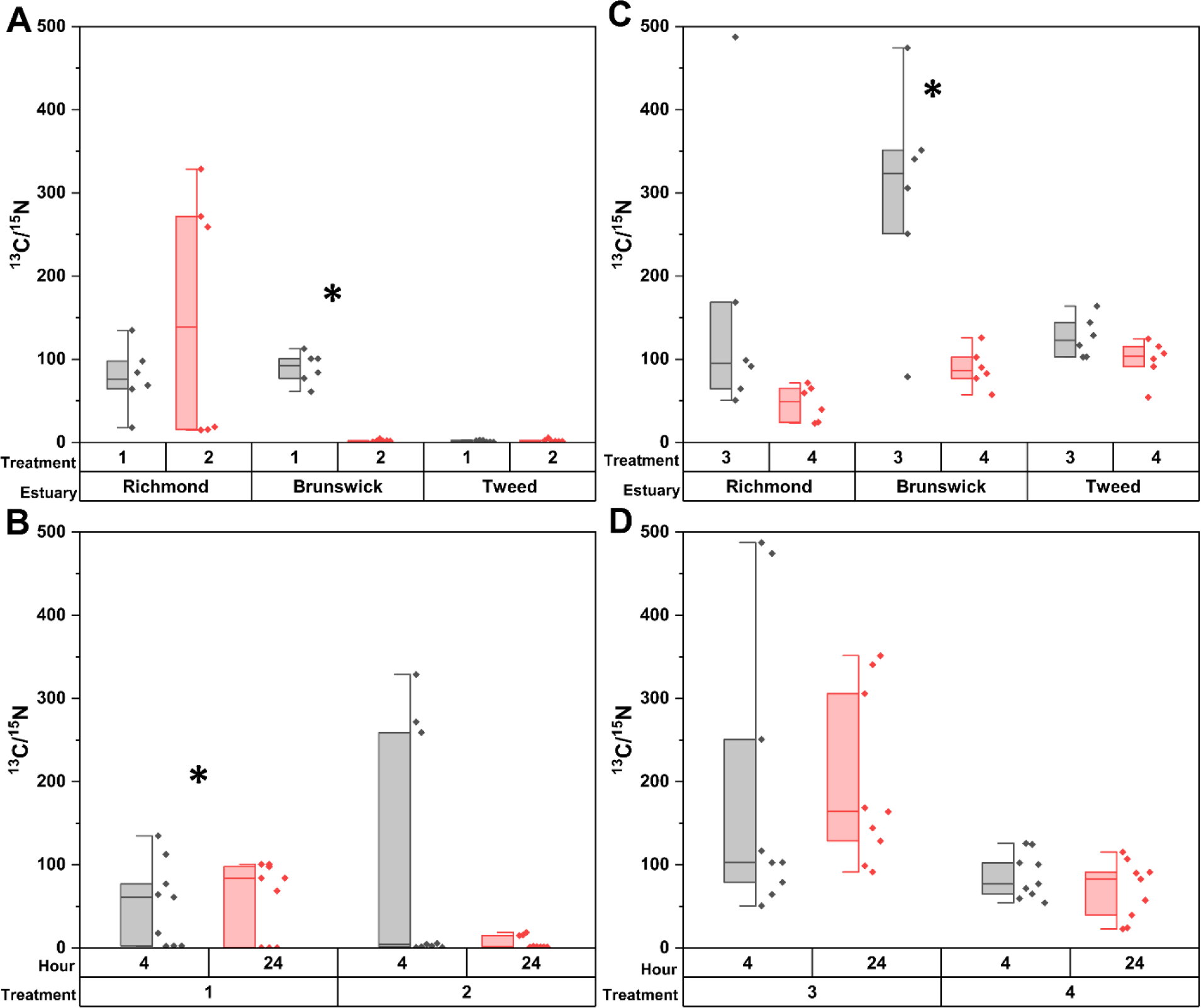
1^3^C to ^15^N ratios retained in sediment from treatment applications applied to intertidal sediment A) within each estuary and B) at 4 and 24 hours across all three estuaries. Dual labelled (^15^N and ^13^C) treatment applications contain: 1) algal dissolved organic matter, 2) an amino acid mixture, 3) glucose and NH_4_^+^, and 4)NaHCO_3_ and NH_4_^+^. Uptake concentrations have been scaled to allow direct comparison between organic (1 and 2) and NH_4_^+^ and glucose or NaHCO_3_ (3 and 4) treatment applications. An asterisk indicates a significant difference (*p*<0.05) between pairings using non-parametric pairwise Wilcoxon sign rank tests.

Bacterial biomass indicated by D/L-Ala was marginally different between treatments (Fig 7A; F_2,71_=2.9, *p*=0.04), but a post hoc Tukey’s test indicated no difference amongst treatments (*p*>0.05; range 76 to 85%). Between estuaries, bacterial biomass was higher in the Richmond than the Tweed (F_2,71_=6.6, *p*<0.01; range 75-86%). Bacterial uptake indicated from the ratio of uptake of ^15^N into the D and L forms of alanine had a significant 3-way interaction that required applying 2-way ANOVAs for Estuary*Treatment to the 4 and 24 h periods separately (Fig. 7B; 4 h, F_5,35_=4.8, *p*<0.01; 24 h, F_5,34_=2.9, *p*=0.03). For the 4 h period, uptake of ^15^N was dominated by bacteria in all treatments except for glucose and NH ^+^ (62-65% vs. 23%, respectively), but by 24 h there was no significant difference between treatments. However, the Tweed estuary had lower bacterial uptake of ^15^N than the other two estuaries (25% vs 53-55%) indicating higher retention of ^15^N in MPB.

**Figure 7:**
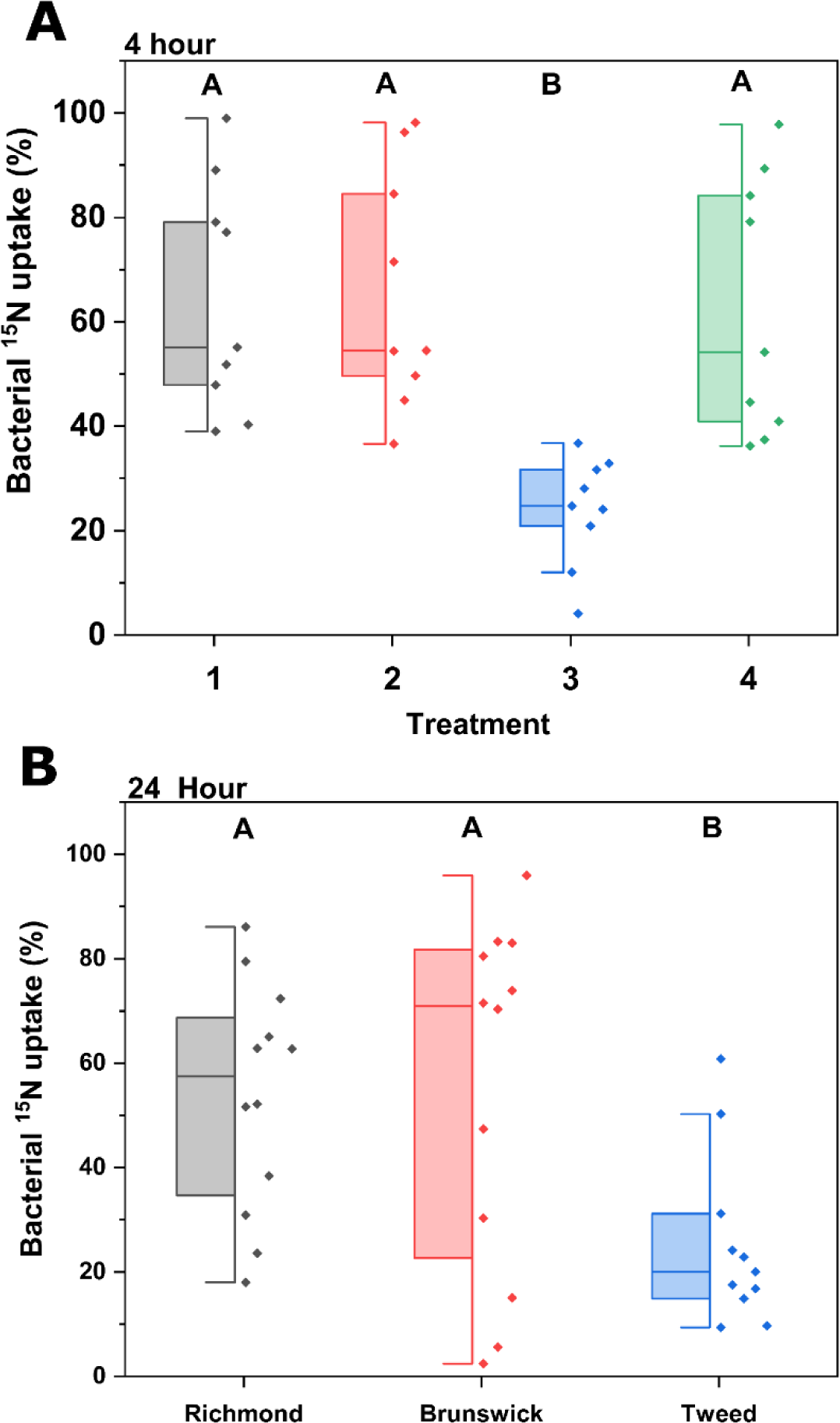
Percentage of total ^15^N incorporated into heterotrophic bacteria within the microbial community. Bacterial uptake is derived from the ^15^N content of D/L-alanine in diatoms and heterotrophic bacteria.

## Discussion

Through the application of dual labelled (^13^C and ^15^N) substrates and mixtures, we have identified the short-term uptake and fate from each organic (algal DOM; treatment 1 and amino acid mixture; treatment 2) and inorganic (NH_4_^+^, glucose; treatment 3 and NH_4_^+^, NaHCO_3_; treatment 4) application to reveal differences in the relative lability of organic materials and the dominant processing pathways during uptake by MPB and heterotrophic bacteria. This work highlights the considerable variability encountered from in situ sediment applications despite the effort taken in targeting sediments from geographically and locally similar settings (e.g. river-dominated muddy sands). This study represents a well-replicated comparison of short-term uptake rates for C and N from multiple substrate types within intertidal sediments, sampling river dominated sub-tropical estuaries across relatively small spatial and temporal scales. Although this work is spatially limited in scope (subtropical East coast of Australia), it helps to clarify the extent of heterogeneity during uptake and short-term processing of C and N across latitudes in similar intertidal settings (Table 1).

Studies using isotope labelling have observed considerable variability between both between replicates and studies (%CV; 5-63% ^13^C; 7-60% ^15^N) in short term uptake and processing of inorganic or organic labeled substrates, regardless of whether the isotope-labelled substrate was applied to slurries or intact sediments (Table 1 and included references). Studies included here represent the bulk of the work undertaken in intertidal mudflats using rare stable isotope labelling techniques for carbon and nitrogen. The variability observed across these 20 studies ranges across orders of magnitude thus often making it difficult to satisfy requirements for statistical analysis, such as homogeneity of variance, even following data despite transformation. Even coefficient of variation is significantly different amongst ^13^C substrate uptake (F_1,24_=53.2, *p*<0.001) and ^15^N substrates (F_1,23_=34.5, *p*=0.044) using a modified signed-likelihood ratio test (Krishnamoorthy and Lee, 2014) on studies with adequate replication for inclusion. Variability does not appear to be a result of climate (e.g. latitude), as variance in ^15^N and ^13^C uptake remains considerable across studies performed in temperate, subtropical and for the single sub-Arctic study (Oakes et al., 2016). Variability during uptake may reflect heterogeneity resulting from the combination of overlying water flow rates, nutrient availability (Høgslund et al., 2023; Riekenberg et al., 2020), sediment composition (Pryshlak et al., 2015; Wallace et al., 2020), substrate availability and quality, and the composition of the microbial community (Carreira et al., 2013). In this study, MPB-C biomass measurements confirm that there was no short-term (24 h) change in biomass resulting from treatment applications between the estuaries examined here. However, despite similar biomass being present across our study, we observed considerable variability in the uptake of labelled substrates. This indicates that the uneven distribution of ^13^C and ^15^N between replicates across other studies may be due to 1) variable diffusion of substrates through porewater or 2) heterogenous distribution of the microbial community that may have caused the development of hotspots of uptake and biogeochemical processing (McClain et al., 2003; Santner et al., 2015).

### Uptake from algal DOM and amino acids

Amino acids can be a large portion of the organic matter pool in marine settings (Cowie and Hedges, 1992). Microbial community uptake of amino acids can vary from minimal, resulting in downward transport of newly fixed autotrophic material via the microbial shunt in the open sea (McCarthy et al., 2004), to complete use by sponge symbionts (Shih et al., 2019). In estuarine sediment, amino acids are expected to be quickly taken up and used by the microbial community as a relatively labile molecule, but use of C and N from these compounds can be either coupled or uncoupled (Veuger and Middelburg, 2007) depending on whether applied amino acids are directly incorporated into biomass (coupled) or degraded into NH_4_^+^ and the accompanying carbon skeleton prior to use by the microbial community. Higher retention of ^13^C from algal DOM than amino acids across the 4 and 24 h periods in the current study (Fig. 2) indicates that ^13^C from DOM was preferentially incorporated into biomass, whereas C from amino acids was more likely to be shunted into supporting respiration and exported from the sediments.

In the organic treatments more ^13^C was taken up from algal DOM than from amino acids (∼150 to 300 µM C m^-2^; Fig. 2A&B), which contrasts with previous work in sediment slurries that observed preferential uptake of carbon from algal-derived amino acids vs urea, a relatively labile substrate (Veuger and Middelburg, 2007). Higher use of ^13^C from algal DOM indicates immediate preferential use of ^13^C vs amino acids (Fig. 2A), likely due to a more complex mixture of ^13^C compounds (e.g. short chain fatty acids, acetate, pyruvate) being incorporated into wider range of biomass rather than being shunted towards supporting metabolism. Uptake was followed with relatively higher retention of ^13^C from algal DOM, a complex mixture of molecules, than from amino acids across all three estuaries (Fig. 2B). Relatively low uptake of ^13^C and higher uptake of ^15^N from amino acids indicates decoupled use between C and N and the dominant uptake pathway for AAs is likely via extracellular amino acid oxidation. Oxidation of amino acids allows for the preferential use of N from AAs and leaves the carbon skeletons as labile DOC to be respired by heterotrophic bacteria and subsequently exported to the water column as DIC (Mulholland et al., 1998). The uptake rates for both organic treatments (Fig. 2) were lower than those observed for glucose (∼ 530 to 2100 µM C m^-2^ Fig. 4), which likely reflects the requirement for algal DOM to be degraded by enzymes before becoming available for uptake.

High ^13^C/^15^N ratios for both organic treatments in the Richmond River and for algal DOM in the Brunswick (Fig. 6A) indicate disproportionally higher immediate uptake and short-term retention of ^13^C from DOM than from the amino acid treatment. Initially high decoupled uptake of ^13^C followed with lower ^13^C/^15^N values at 24 h in the Brunswick reflect immediate use of labile carbon. These C substrates are potentially carbohydrates and sugars included in the algal DOM treatment and carbon skeletons from amino acids to support immediate respiration. Similarly high ^13^C values at 24 h across all three estuaries (Fig. 2) for algal DOM indicate considerable short-term retention of the newly fixed ^13^C observed at 4 h. Higher uptake and retention likely indicates incorporation of newly fixed carbon into microbial biomass or refractory molecules within sediment. ^13^C/^15^N ratios for algal DOM were lower than the applied amino acid treatment (4.4) in the Brunswick and Tweed estuaries (2.2±1.5; Fig. 6A). Higher uptake of N from amino acids indicates decoupled initial uptake of ^15^N and ^13^C reflecting higher rates of microbial N use followed with preferential retention of ^15^N within either biomass or molecules in the bulk sediment. The presence of excess ^15^N in D-alanine at 4 h (Fig. 7A) confirms that uptake from both organic treatments was dominated by heterotrophic bacteria. Differences in substrate use between estuaries reflect the balance between selective uptake into biomass due to substrate quality, respiration, and remineralization in the short-term.

#### Uptake from glucose, bicarbonate, and ammonium

Between the treatments comparing NH_4_^+^ use in the Brunswick and Tweed estuaries, considerably more ^13^C was taken up from the glucose and NH_4_^+^ treatment than was newly fixed into MPB-C from the NaHCO_3_^-^ and NH_4_^+^ treatments (Fig. 4A). Tight coupling in the microbial community is expected in conditions where carbon is limited to production by MPB under nutrient limited conditions (Riekenberg et al., 2020) and glucose is therefore the predominant form of newly fixed carbon (Moerdijk-Poortvliet et al., 2018) with limited alternative sources, especially during sediment exposure. In the Richmond River estuary ^13^C uptake was similar between the two treatments indicating similar use of newly fixed MPB-C and labile sugars with uptake happening in a comparable range to the Tweed. The Richmond River has been previously observed to be carbon limited (Oakes and Eyre, 2014), so it is not surprising that there was similar uptake of ^13^C between the two treatments, indicating quick uptake and recycling of labeled substrates by MPB and heterotrophic bacteria regardless of substrate form. This suggests comparable use of carbon between newly fixed MPB-C and glucose which would be expected from a microbial community with tight coupling between MPB and heterotrophic bacteria, with heterotrophic bacteria processing extracellular polymeric substances and producing NH_4_^+^ that then supports MPB (Cook et al., 2007).

In contrast, in the Brunswick and Tweed, higher uptake by the microbial community of ^13^C from glucose than from newly fixed MPB-C indicates stimulated uptake of carbon in excess of what occurs during production of MPB-C. This may be a result of the bacterial community using glucose to support alternative processing pathways such as denitrification (Morelle et al., 2022), but this possibility remains speculative without inclusion of a direct measurement of N_2_ fluxes in this study. Despite this limitation, these values serve as an *in situ* comparison between heterotrophic and autotrophic C uptake and processing. Previous work incorporating direct measurements of N_2_ fluxes have identified contributions from denitrification to the nitrogen budget in the Richmond (∼3%; Riekenberg et al., 2020) and higher contributions in the Brunswick (∼21%; Eyre et al., 2016a). The similar uptake of ^15^N from observed for both NH_4_ ^+^ treatments in the Brunswick may be due to export of ^15^N via coupled nitrification-denitrification for treatment 3 as evident from the disproportionately high uptake of ^13^C from glucose and increased incorporation of ^15^N in treatment 4 (NaHCO_3_, Fig. 7A). The comparable uptake of ^15^N between treatments is deceptive since application rates for NH ^+^ were ∼3× higher in the glucose treatment. Low uptake and retention of ^15^N could potentially indicate either considerably higher export rates or a low threshold for ^15^N saturation in these systems, but saturation is unlikely due to higher uptake of ^15^N found in the amino acid treatment. Decreased uptake and retention of ^15^N potentially indicates export from heterotrophic bacteria likely via coupled nitrification-denitrification. Processing along this pathway leaves more of the total ^15^N contained in MPB, effectively leaving MPB as a N reservoir in situations where denitrifying bacteria have been stimulated, but again, this remains speculative without measured N_2_ efflux between the treatments.

Apart from stimulated uptake of ^13^C from glucose in the Brunswick, ^13^C/^15^N ratios for treatments 3 and 4 were equivalent across estuaries and time periods (Fig. 6C & D). This similarity indicates that ^13^C uptake and processing occurs at similar rates regardless of the form of C that the heterotrophs encounter, either simple sugars or newly fixed MPB-C. Despite the similarity in uptake rates, the variability between replicates was considerably higher for the glucose application largely driven by variable ^13^C uptake (Fig, 4). This suggests that hotspots (McClain et al., 2003) for simple sugar uptake and processing occur across all of the sediments examined, where heterotrophic bacteria have limited access to labile carbon but have the ability to readily shift if labile substrates become available episodically. High variability in uptake occurs across sediments on the cm scale across all the intertidal sediment examined here and likely reflects a flexible life strategy for a microbial community that experiences routine variations in substrate availability. This flexibility in supporting metabolism is well suited to intertidal environments where tidal inundation and exposure occur and substrates are likely to be briefly available and then promptly limiting depending on the tidal cycle, or in estuarine settings where rainfall can quickly change inputs to the system.

#### Generalities between organic and inorganic processing

Carbon uptake and processing is more variable than expected and ranged from almost complete exclusion in the case of amino acids to glucose uptake as high as 3.5 mg m^-2^ in a single replicate in the Brunswick estuary. The wide ranges and considerable variability throughout this study for the uptake of ^13^C highlight both the varied pathways that support for MPB and heterotrophs and how tightly coupled MPB production and heterotrophic processing are, depending on substrate availability. Direct stimulation of heterotrophs with the addition of a labile sugar results in variable uptake of ^13^C and lower ^15^N uptake and retention considering the 3× higher NH_4_ ^+^ application between treatments 3 and 4. ^13^C uptake was highly variable between replicates amongst the glucose applications and suggests that hotspots for processing quickly develop in areas where heterotrophic communities using the coupled nitrification-dentrification pathway quickly make use of the added substrates. In contrast, uptake and processing via carbon fixation as MPB-C was lower and less variable across the estuaries indicating more homogeneous incorporation and processing between MPB and heterotrophic bacterial communities. It is unsurprising that MPB-C fixation is the dominant pathway for carbon support of the microbial community in photic intertidal sediments, and this is therefore reflected in the wider data set. Similarly, the relatively lower but consistent uptake of ^13^C from algal DOM implies that this substrate mixture was less available than newly fixed MPB-C but was still readily incorporated and processed via heterotrophy. In contrast, amino acid C was generally preferentially excluded with the notable exception of 3 replicates in the Richmond estuary, indicating the dominant retention of N from these labile substrates, except where carbon limitation necessitates the uptake of amino acid C. It is unclear from the current work what C limitation threshold prompts the switch to amino acid C use rather than exclusion or complete export of amino acid C via respiration. However, the limited use of amino acid C nevertheless implies an energetic disadvantage to processing this material, suggesting that additional steps (enzymatic or membrane transfer) are required to make amino acid C energetically profitable to support respiration.

#### Conclusion

This work used well replicated applications of organic and inorganic substrates across three estuaries to explore 1) the relative uptake and processing between amino acids and a relatively less labile algal DOM mixture and 2) the processing of labile sugars versus newly fixed MPB-C using two applications of ^15^NH_4_^+^ coupled with glucose and NaH^13^CO_3_^+^. In contrast to our expectations, amino acid ^13^C was generally excluded, although not completely, indicating the preferential use of N from these compounds except in rare situations of exceptional C limitation. ^15^N uptake from amino acids was higher than from algal DOM and reflects the relatively more refractory nature of the algal DOM mixture and supporting our hypothesis of labile materials being more readily used, but both substrates demonstrated clear use and retention across 24 h. Uptake and processing of ^13^C from glucose (dominated by heterotrophic bacteria) was considerably more variable and somewhat higher than for ^13^C from NaHCO_3_^+^ (dominated by MPB) which likely indicates increased processing from hotspots of coupled nitrification-denitrification occurring across the estuarine sediments. However, this result remains speculative due to not measuring N_2_ fluxes simultaneously in this study, despite observing comparable ^15^N uptake and retention for NH_4_ between applications that were scaled 3 times higher for the glucose treatment. In contrast to our expectations, uptake and processing routed through MPB via fixation pathways was generally lower for ^13^C, but variable for ^15^N across estuaries indicating that local processes likely contribute to regulating the rates of fixation and processing MPB-derived organic matter even amongst similar river-dominated intertidal sediments.

#### Implications

The considerable variability in substrate use and processing observed in this study highlights a major issue in the investigation of process rates in estuarine samplings. Hotspots and hot moments, localized areas or times of disproportionately high biogeochemical processing, are recognized to routinely develop and persist in sediments largely regulated by substrate availability (McClain et al., 2003) and contribute to the considerable variability observed in uptake and processing within labelling studies. This processing couples carbon and nitrogen use, but the majority of the studies examining denitrification only characterize the processing of N, and not C (Douglas et al., 2022; Peng et al., 2022). Coupled examination of both C and N can help identify which substrates are available and how substrate quality changes affect the microbial community composition and processing pathways. This work further highlights the need for in situ studies coupling gas measurements to treatment applications to ensure that gas flux rates for quantification of target biogeochemical pathways are possible. MPB-dominated sediments can display a wide range of uptake and processing rates resulting with considerable variability (Table 1) that is difficult to adequately investigate using traditional statistical techniques. High variability between replicates is characteristic of substrate processing in sediments, and lowered labeling concentrations that better reflect environmental conditions within in situ applications appear to exacerbate this problem. Future work should aim for sufficient replication to allow for statistical analysis while accounting for limitations of labour and cost associated with laboratory analysis, while also acknowledging that replicate variability may very well still interfere with data analysis. Further work, development, and application of analytical techniques used for data from systems that tend to be episodic and demonstrate high variability would help to further separate and identify signals from biogeochemical processing (i.e., hot spots) from the noise (replicate variability).

## Supporting information

Supporting_info_uptake_MS

## Acknowledgements

We would like to thank J. Ossebaar, R. van Bommel, and J.L. Riekenberg for laboratory support in processing and analyzing the samples supporting this work. J.L. Riekenberg also contributed to fieldwork in the hectic time between PhD completion and an international move, which is greatly appreciated.

