## Supporting_info_uptake_MS for "Hot spots drive uptake and short-term processing of organic and inorganic carbon and nitrogen in intertidal sediments"

Supplemental Figure 1: Labelling dosages for both $^{13}C$ and $^{15}N$ for all treatment applications.

| | **μmol $L^{-1}$ C** | **μmol $L^{-1}$ N** | **vol (L)** | **$m^2$** | **μmol C $m^{-2}$** | **μmol N $m^{-2}$** | **μmol $^{13}C$ $m^{-2}$** | **adjustment factor** | **μmol $^{15}N$ $m^{-2}$** | **adjustment factor** |
|---|---|---|---|---|---|---|---|---|---|---|
| **Algal DON** | 162 | 60 | 0.02 | 0.32 | **10.1** | **3.8** | 0.5 | X | 1.8 | 1.4 |
| **Amino acids** | 211 | 60 | 0.02 | 0.32 | **13.2** | **3.8** | 13.1 | 25.8 | 3.7 | 3.0 |
| **Glucose + $NH_4^+$** | 1100 | 60 | 0.02 | 0.32 | **68.8** | **3.8** | 68.1 | X | 3.7 | 3.0 |
| **$NaHCO_3$ + $NH_4^+$** | 1800 | 20 | 0.02 | 0.32 | **112.5** | **1.3** | 110.3 | 1.6 | 1.2 | X |

Table S1: Statistical tests supporting between estuary treatment applications

| Wilcoxon Sign ranked tests | Estuary | treatment | Comparison | Z | p |
|---|---|---|---|---|---|
| $^{15}N$ | Richmond | Algal DOM | 4 vs 24 h | 0.3 | 0.2 |
| | Richmond | Amino acid | 4 vs 24 h | -0.3 | 0.8 |
| | Richmond | Glucose and $NH_4^+$ | 4 vs 24 h | 0 | 1 |
| | Richmond | $NaHCO_3$ and $NH_4^+$ | 4 vs 24 h | -1.3 | 0.2 |
| | Tweed | Algal DOM | 4 vs 24 h | -1.3 | 0.2 |
| | Tweed | Amino acid | 4 vs 24 h | 0 | 1 |
| | Tweed | Glucose and $NH_4^+$ | 4 vs 24 h | 0.8 | 0.4 |
| | Tweed | $NaHCO_3$ and $NH_4^+$ | 4 vs 24 h | 0 | 1 |
| | Brunswick | Algal DOM | 4 vs 24 h | -1.3 | 0.2 |
| | Brunswick | Amino acid | 4 vs 24 h | -1.3 | 0.2 |
| | Brunswick | Glucose and $NH_4^+$ | 4 vs 24 h | 1.3 | 0.2 |
| | Brunswick | $NaHCO_3$ and $NH_4^+$ | 4 vs 24 h | 0.3 | 0.8 |

| Wilcoxon Sign ranked tests | Estuary | treatment | Comparison | Z | p |
|---|---|---|---|---|---|
| $^{13}C$ | Richmond | Algal DOM | 4 vs 24 h | 0.8 | 0.4 |
| | Richmond | Amino acid | 4 vs 24 h | 1.3 | 0.2 |
| | Richmond | Glucose and $NH_4^+$ | 4 vs 24 h | 0 | 1 |
| | Richmond | $NaHCO_3$ and $NH_4^+$ | 4 vs 24 h | 1.3 | 0.2 |
| | Tweed | Algal DOM | 4 vs 24 h | -0.8 | 0.4 |
| | Tweed | Amino acid | 4 vs 24 h | 0.3 | 0.8 |
| | Tweed | Glucose and $NH_4^+$ | 4 vs 24 h | 0 | 1 |
| | Tweed | $NaHCO_3$ and $NH_4^+$ | 4 vs 24 h | 1.3 | 0.2 |
| | Brunswick | Algal DOM | 4 vs 24 h | 0.8 | 0.4 |
| | Brunswick | Amino acid | 4 vs 24 h | 0.3 | 0.8 |
| | Brunswick | Glucose and $NH_4^+$ | 4 vs 24 h | -0.8 | 0.4 |
| | Brunswick | $NaHCO_3$ and $NH_4^+$ | 4 vs 24 h | 0 | 1 |

| Wilcoxon Sign ranked tests | Estuary | treatment | Comparison | Z | p |
|---|---|---|---|---|---|
| D/L-Ala | Richmond | Algal DOM | 4 vs 24 h | 0.8 | 0.4 |
| | Richmond | Amino acid | 4 vs 24 h | 0.3 | 0.8 |
| | Richmond | Glucose and $NH_4^+$ | 4 vs 24 h | 0.3 | 0.2 |
| | Richmond | NAHCO3 and $NH_4^+$ | 4 vs 24 h | 1 | 1 |
| | Tweed | Algal DOM | 4 vs 24 h | 1 | 1 |
| | Tweed | Amino acid | 4 vs 24 h | -0.8 | 0.4 |
| | Tweed | Glucose and $NH_4^+$ | 4 vs 24 h | 1 | 1 |
| | Tweed | NAHCO3 and $NH_4^+$ | 4 vs 24 h | 1.3 | 0.2 |
| | Brunswick | Algal DOM | 4 vs 24 h | -1.3 | 0.2 |
| | Brunswick | Amino acid | 4 vs 24 h | 1.3 | 0.2 |
| | Brunswick | Glucose and $NH_4^+$ | 4 vs 24 h | 1.3 | 0.2 |
| | Brunswick | $NaHCO_3$ and $NH_4^+$ | 4 vs 24 h | 0.9 | 0.4 |

| Wilcoxon Sign ranked tests | Estuary | Treatment | Z | p |
|---|---|---|---|---|
| $^{15}N$ | Richmond | Algal DOM vs Amino acids | -2.1 | 0.036 |
| | Richmond | Glucose and $NH_4^+$ vs $NaHCO_3^-$ and $NH_4^+$ | -2.1 | 0.036 |
| | Tweed | Algal DOM vs Amino acids | -2.1 | 0.036 |
| | Tweed | Glucose and $NH_4^+$ vs $NaHCO_3^-$ and $NH_4^+$ | -1.3 | 0.2 |
| | Brunswick | Algal DOM vs Amino acids | -2.1 | 0.036 |
| | Brunswick | Glucose and $NH_4^+$ vs $NaHCO_3^-$ and $NH_4^+$ | 2.1 | 0.036 |

| Wilcoxon Sign ranked tests | Estuary | Treatment | Z | P |
|---|---|---|---|---|
| $^{13}C$ | Richmond | Algal DOM vs Amino acids | 2.1 | 0.036 |
| | Richmond | Glucose and $NH_4^+$vs $NaHCO_3^-$ and $NH_4^+$ | 1.1 | 0.3 |
| | Tweed | Algal DOM vs Amino acids | 2.1 | 0.036 |
| | Tweed | Glucose and $NH_4^+$vs $NaHCO_3^-$ and $NH_4^+$ | 1.9 | 0.06 |
| | Brunswick | Algal DOM vs Amino acids | 2.1 | 0.036 |
| | Brunswick | Glucose and $NH_4^+$vs $NaHCO_3^-$ and $NH_4^+$ | 2.1 | 0.036 |

| Wilcoxon Sign ranked tests | Estuary | Treatment | Z | p |
|---|---|---|---|---|
| $^{13}C/^{15}N$ | Richmond | Algal DOM vs Amino acids | -0.8 | 0.4 |
| | Richmond | Glucose and $NH_4^+$ vs $NaHCO_3^-$ and $NH_4^+$ | 1.5 | 0.1 |
| | Tweed | Algal DOM vs Amino acids | -1.1 | 0.3 |
| | Tweed | Glucose and $NH_4^+$vs $NaHCO_3^-$ and $NH_4^+$ | 1.5 | 0.1 |
| | Brunswick | Algal DOM vs Amino acids | 2.1 | 0.04 |
| | Brunswick | Glucose and $NH_4^+$vs $NaHCO_3^-$ and $NH_4^+$ | 2.1 | 0.04 |

| Wilcoxon Sign ranked tests | Estuary | Treatment | Z | p |
|---|---|---|---|---|
| D/L-Ala | Richmond | Algal DOM vs Amino acids | 0.8 | 0.4 |
| | Richmond | Glucose and $NH_4^+$vs $NaHCO_3^-$ and $NH_4^+$ | -1 | 0.3 |
| | Tweed | Algal DOM vs Amino acids | 0 | 1 |
| | Tweed | Glucose and $NH_4^+$vs $NaHCO_3^-$ and $NH_4^+$ | -1.9 | 0.06 |
| | Brunswick | Algal DOM vs Amino acids | -0.4 | 0.7 |
| | Brunswick | Glucose and $NH_4^+$vs $NaHCO_3^-$ and $NH_4^+$ | -0.5 | 0.6 |

Table S2 Statistical tests supporting across estuary temporal comparisons

| Wilcoxon Sign ranked tests | Hour | Treatment | Z | p |
|---|---|---|---|---|
| Pooling treatments | 4 | Algal DOM vs Amino acids | -2.6 | 0.00915 |
| $^{15}N$ | 24 | Algal DOM vs Amino acids | -2.6 | 0.00915 |
| | 4 | Glucose and $NH_4^+$ vs $NaHCO_3^-$ and $NH_4^+$ | -0.2 | 0.8 |
| | 24 | Glucose and $NH_4^+$ vs $NaHCO_3^-$ and $NH_4^+$ | -1.7 | 0.1 |

| Hour | Treatment | Z | p |
|---|---|---|---|
| 4 vs 24 | Algal DOM | -2.6 | 0.00915 |
| 4 vs 24 | Amino acids | -1.1 | 0.3 |
| 4 vs 24 | Glucose and $NH_4^+$ | 1.4 | 0.2 |
| 4 vs 24 | $NaHCO_3^-$ and $NH_4^+$ | -0.6 | 0.6 |

| Wilcoxon Sign ranked tests | Hour | Treatment | Z | p |
|---|---|---|---|---|
| Pooling treatments | 4 | Algal DOM vs Amino acids | 2.6 | 0.00915 |
| $^{13}C$ | 24 | Algal DOM vs Amino acids | 2.6 | 0.00915 |
| | 4 | Glucose and $NH_4^+$ vs $NaHCO_3^-$ and $NH_4^+$ | 1.7 | 0.1 |
| | 24 | Glucose and $NH_4^+$ vs $NaHCO_3^-$ and $NH_4^+$ | 2.6 | 0.00915 |

| | Hour | Treatment | Z | p |
|---|---|---|---|---|
| | 4 vs 24 | Algal DOM | 0.4 | 0.7 |
| | 4 vs 24 | Amino acids | 1.4 | 0.2 |
| | 4 vs 24 | Glucose and $NH_4^+$ | -0.6 | 0.6 |
| | 4 vs 24 | $NaHCO_3^-$ and $NH_4^+$ | 1.9 | 0.06 |

| Wilcoxon Sign ranked tests | Hour | Treatment | Z | p |
|---|---|---|---|---|
| Pooling treatments | 4 | Algal DOM vs Amino acids | 2.6 | 0.009 |
| $^{13}C/^{15}N$ | 24 | Algal DOM vs Amino acids | 2.6 | 0.009 |
| | 4 | Glucose and $NH_4^+$ vs $NaHCO_3^-$ and $NH_4^+$ | 1.2 | 0.2 |
| | 24 | Glucose and $NH_4^+$ vs $NaHCO_3^-$ and $NH_4^+$ | 2.6 | 0.004 |

| | Hour | Treatment | Z | p |
|---|---|---|---|---|
| | 4 vs 24 | Algal DOM | 2.1 | 0.03 |
| | 4 vs 24 | Amino acids | 1 | 0.3 |
| | 4 vs 24 | Glucose and $NH_4^+$ | -0.7 | 0.5 |
| | 4 vs 24 | $NaHCO_3^-$ and $NH_4^+$ | 1.4 | 0.2 |

| Wilcoxon Sign ranked tests | Hour | Treatment | Z | p |
|---|---|---|---|---|
| Pooling treatments | 4 vs 24 | Algal DOM | 0.6 | 0.6 |
| D/L-Ala | 4 vs 24 | Amino acids | 1.2 | 0.2 |
| | 4 vs 24 | Glucose and $NH_4^+$ | -0.2 | 0.8 |
| | 4 vs 24 | NAHCO3- and $NH_4^+$ | 1.1 | 0.29 |

Table S3: ANOVAS supporting estuary, time, and treatment applications for D/L-alanine

Levene's D/L-Ala $F_{3,67}=1.52$ $p=0.2$

3-way ANOVA D/L-Ala

| | DF | Sum of Squares | Mean Square | F Value | P Value | |
|---|---|---|---|---|---|---|
| Est | 2 | 2753.293 | 1376.646 | 3.08258 | 0.05522 | + |
| TRT | 3 | 11428 | 3809.334 | 8.52984 | 1.25E-04 | * |
| Hour | 1 | 1684.216 | 1684.216 | 3.77129 | 0.05815 | + |
| Est * TRT | 6 | 4407.373 | 734.5621 | 1.64483 | 0.15594 | |
| Est * Hour | 2 | 4350.04 | 2175.02 | 4.87029 | 0.01196 | * |
| TRT * Hour | 3 | 1936.184 | 645.3948 | 1.44516 | 0.24162 | |
| Est * TRT * Hour | 6 | 6162.578 | 1027.096 | 2.29987 | 0.04992 | * |
| Model | 23 | 31872.86 | 1385.776 | 3.10302 | 4.97E-04 | * |
| Error | 47 | 20989.69 | 446.5892 | | | |
| Corrected Total | 70 | 52862.55 | | | | |

2-Way ANOVA
Est*TRT for 4 h

| | DF | Sum of Squares | Mean Square | F Value | P Value | |
|---|---|---|---|---|---|---|
| Estuary | 2 | 109.226 | 54.613 | 0.11928 | 0.88798 | |
| Treatment | 3 | 10900.39 | 3633.463 | 7.93594 | 4.84E-04 | * |
| Model | 5 | 11091.34 | 2218.268 | 4.84497 | 0.00229 | * |
| Error | 30 | 13735.48 | 457.8493 | | | |
| Corrected Total | 35 | 24826.82 | | | | |

2-Way ANOVA
Est*TRT for 24 h

| | DF | Sum of Squares | Mean Square | F Value | P Value | |
|---|---|---|---|---|---|---|
| Estuary | 2 | 6227.918 | 3113.959 | 5.10028 | 0.01265 | * |
| Treatment | 3 | 2530.786 | 843.5955 | 1.38171 | 0.26807 | |
| Model | 5 | 8903.502 | 1780.7 | 2.91657 | 0.02979 | * |
| Error | 29 | 17705.84 | 610.5461 | | | |
| Corrected Total | 34 | 26609.34 | | | | |

Levene's HOV C Biomass F2,69=1.8 p=0.2

3-way ANOVA C Biomass

| | DF | Sum of Squares | Mean Square | F Value | P Value | |
|---|---|---|---|---|---|---|
| Estuary | 2 | 1436.645 | 718.3224 | 9.10903 | 4.43E-04 | * |
| Treatment | 3 | 749.0921 | 249.6974 | 3.16641 | 0.03272 | * |
| Hour | 1 | 201.3815 | 201.3815 | 2.55372 | 0.1166 | |
| Estuary * Treatment | 6 | 1508.584 | 251.4306 | 3.18839 | 0.01027 | * |
| Estuary * Time | 2 | 375.7021 | 187.8511 | 2.38214 | 0.10319 | |
| Treatment * Time | 3 | 65.92911 | 21.97637 | 0.27868 | 0.8405 | |
| Estuary * Treatment * Time | 6 | 953.2705 | 158.8784 | 2.01473 | 0.08191 | |
| Model | 23 | 5347.199 | 232.4869 | 2.94816 | 7.88E-04 | * |
| Error | 48 | 3785.196 | 78.85826 | | | |
| Corrected Total | 71 | 9132.395 | | | | |

2-Way ANOVA

| Est*Trt | DF | Sum of Squares | Mean Square | F Value | P Value | |
|---|---|---|---|---|---|---|
| Estuary | 2 | 1373.848 | 686.9238 | 6.62545 | 0.00239 | * |
| Treatment | 3 | 885.5482 | 295.1827 | 2.84707 | 0.04415 | * |
| Model | 5 | 2289.543 | 457.9086 | 4.41657 | 0.00157 | * |
| Error | 66 | 6842.852 | 103.6796 | | | |
| Corrected Total | 71 | 9132.395 | | | | |

Table S3: CV comparison across C and N treatments in this study

CV between sites

| C Treatments | | N | mean | CV | test statistic | *p* | |
|---|---|---|---|---|---|---|---|
| | 1 | 3 | 16.2 | 0.438272 | 19.4 | 0.0465 | * |
| | 1 | 3 | 52 | 0.625 | | | |
| | 1 | 3 | 15.1 | 0.145695 | | | |
| | 2 | 3 | 196 | 0.591837 | | | |
| | 2 | 3 | 295 | 0.049831 | | | |
| | 2 | 3 | 306 | 0.074183 | | | |
| | 3 | 3 | 765 | 0.797386 | | | |
| | 3 | 3 | 2166 | 0.614035 | | | |
| | 3 | 3 | 824 | 0.151699 | | | |
| | 4 | 3 | 664 | 0.103916 | | | |
| | 4 | 3 | 974 | 0.243326 | | | |
| | 4 | 3 | 417 | 0.347722 | | | |

| N Treatments | | | | | test statistic | *p* |
|---|---|---|---|---|---|---|
| | 1 | 3 | 16.2 | 0.438272 | 13 | 0.29 |
| | 1 | 3 | 52 | 0.625 | | |
| | 1 | 3 | 15.1 | 0.145695 | | |
| | 2 | 3 | 196 | 0.591837 | | |
| | 2 | 3 | 295 | 0.049831 | | |
| | 2 | 3 | 306 | 0.074183 | | |
| | 3 | 3 | 765 | 0.797386 | | |
| | 3 | 3 | 2166 | 0.614035 | | |
| | 3 | 3 | 824 | 0.151699 | | |
| | 4 | 3 | 664 | 0.103916 | | |
| | 4 | 3 | 974 | 0.243326 | | |
| | 4 | 3 | 417 | 0.347722 | | |

Table S4: CV comparison across C and N substrate applications across studies contained in Table 1

| C Treatments | | mean | CV | | test statistic | *p* | |
|---|---|---|---|---|---|---|---|
| | 1 | 16651 | 0.353612 | 4 | 53.2 | <0.001 | * |
| | 2 | 11250 | 0.048 | 2 | | | |
| | 3 | 1190 | 0.05042 | 2 | | | |
| | 4 | 2770 | 0.185921 | 3 | | | |
| | 5 | 1549 | 0.090381 | 3 | | | |
| | 6 | 892 | 0.035874 | 2 | | | |
| | 7 | 1809 | 0.060807 | 2 | | | |
| | 8 | 12000 | 0.375 | 2 | | | |
| | 9 | 7700 | 0.025974 | 3 | | | |
| | 10 | 664 | 0.103916 | 3 | | | |
| | 11 | 974 | 0.243326 | 3 | | | |
| | 12 | 417 | 0.347722 | 3 | | | |
| | 13 | 765 | 0.797386 | 3 | | | |
| | 14 | 2166 | 0.614035 | 3 | | | |
| | 15 | 824 | 0.151699 | 3 | | | |
| | 16 | 259 | 0.335907 | 3 | | | |
| | 17 | 196 | 0.591837 | 3 | | | |
| | 18 | 295 | 0.049831 | 3 | | | |
| | 19 | 306 | 0.074183 | 3 | | | |
| | 20 | 16.2 | 0.438272 | 3 | | | |
| | 21 | 52 | 0.625 | 3 | | | |
| | 22 | 15.1 | 0.145695 | 3 | | | |
| | 23 | 1645 | 0.064438 | 3 | | | |
| | 24 | 526 | 0.076046 | 2 | | | |

CV between studies

| N Treatments | mean | CV | n | test statistic | *p* | |
|---|---|---|---|---|---|---|
| 1 | 10.2 | 0.107843 | 3 | 34.5 | 0.0442 | * |
| 2 | 9.6 | 0.072917 | 3 | | | |
| 3 | 4.8 | 0.333333 | 3 | | | |
| 4 | 5.8 | 0.448276 | 3 | | | |
| 5 | 9.3 | 0.451613 | 3 | | | |
| 6 | 7.8 | 0.217949 | 3 | | | |
| 7 | 88.4 | 0.475113 | 2 | | | |
| 8 | 3.4 | 0.441176 | 3 | | | |
| 9 | 3.6 | 0.222222 | 3 | | | |
| 10 | 1.1 | 0.090909 | 3 | | | |
| 11 | 9.8 | 0.418367 | 3 | | | |
| 12 | 30.5 | 0.586885 | 3 | | | |
| 13 | 6.2 | 0.145161 | 3 | | | |
| 14 | 1148 | 0.186411 | 3 | | | |
| 15 | 665 | 0.273684 | 3 | | | |
| 16 | 276 | 0.373188 | 3 | | | |
| 17 | 216.7 | 0.20766 | 2 | | | |
| 18 | 124 | 0.183871 | 2 | | | |
| 19 | 737 | 0.183175 | 6 | | | |
| 20 | 245 | 0.065306 | 6 | | | |
| 21 | 124 | 0.184 | 6 | | | |
| 22 | 737 | 0.183 | 6 | | | |
| 23 | 245 | 0.065 | 2 | | | |
